# Structure of IgG-Fc hexamer reveals a mutual lock-and-key mode of Fc-Fc interaction

**DOI:** 10.1101/2022.02.24.481884

**Authors:** Daopeng Yuan, Shuaixiang Zhou, Haiqing Ni, Weixin Yang, Xiao Fang, Yarong Gao, Zhiyuan Shao, Dongmei Bai, Zhihai Wu, Jia Zou, Lu Liu, Jiahui Shi, Nan Zheng, Michael Yu, Yongjun Liu, Xiaodong Xiao, Bingliang Chen, Changshou Gao

## Abstract

IgG exists mainly as monomer, but recent studies suggest that IgG forms hexamer to mediate antibody functions. Although many structures of IgG monomer and its fragments are determined, there is no IgG hexamer structure at atomic level due to the weak Fc-Fc interactions. Here we engineered a hexameric IgG with IgM tailpiece fusion and determined the structure by cryo-EM. IgG-Fc hexamer forms hexagon symmetry with the six Fcs lie in a plane with Fc-Fc interaction in a mutual lock-and-key mode. This structure provides structural insights into Fc-Fc interaction of IgG and reveals molecular basis for its function.

## Introduction

Antibodies are the most important therapeutic proteins with more than 100 approved by FDA^1^. There are five major classes of immunoglobulins in human body, IgG, IgD, IgE, IgM and IgA. IgG, IgD and IgE exist mainly as monomers, whereas IgM and IgA function mainly as polymers mediated by tailpiece extension of Fc^2^. IgG is the predominant isotype in the human body and widely used in therapy due to its Fc function, such as antibody-dependent cellular cytotoxicity (ADCC), antibody-dependent cellular phagocytosis (ADCP) and complement dependent cytotoxicity (CDC).

Fc-Fc interactions of IgG molecules are important in the formations of immune complex and complement activation^3^. Earlier structural studies of IgG showed that IgG-Fc formed a hexamer due to crystal packing^4^. Low resolution structure of IgG-C1 complex and biophysical study showed that IgG can form hexamer on antigen-coated liposome surface via Fc-Fc interaction to activate complement pathway^5–7^. However, there are no structures of hexameric IgG at atomic level to show its assembly.

IgM tailpiece has been fused to IgG and include a L309C (EU numbering) point mutation to form hexameric IgG^8^. IgM-Fc structures were determined by cryo-EM to gain insights into the assembly of IgM and molecular mechanisms of polymerization^9,10^. Based on the structures of IgG and IgM, we designed a IgG hexamer with fusion of IgM tailpiece and incorporated of two mutants at tailpiece to improve polymerization ability. Here we also determined the structure of hexameric IgG-Fc to reveal its assembly in a mutual lock-and-key mode for the first time.

### Engineering of Hexameric IgG

IgG crystal packing formed IgG-Fc hexamer (Fig. 1a, PDB code 1HZH) was superimposed to IgM-Fc cryo-EM structure (Fig. 1b, PDB code 6KXS). The overlapped structure (Fig. 1c) shows that 5 copies of IgG-Fc fit well with 5 IgM-Fcs, with the J-chain occupying the site of a sixth IgG-Fc. Of particular interest, the C-terminus of IgM-Fc and IgG-Fc have different orientations due to the difference of P445 in IgG-Fc and corresponding residue T556 in IgM-Fc (Fig. 1d). In order to make a better fusion of IgM tailpiece to IgG-Fc, we swapped IgG-Fc C-terminal residues S442-K447 to corresponding residues D553-K558 of IgM-Fc (Fig. 1e). Two point mutations in the tailpiece, V567I and A572G, were combined with L309C in IgG-Fc to improve polymerization ability (Supplementary Fig. 1).

**Fig.1.**
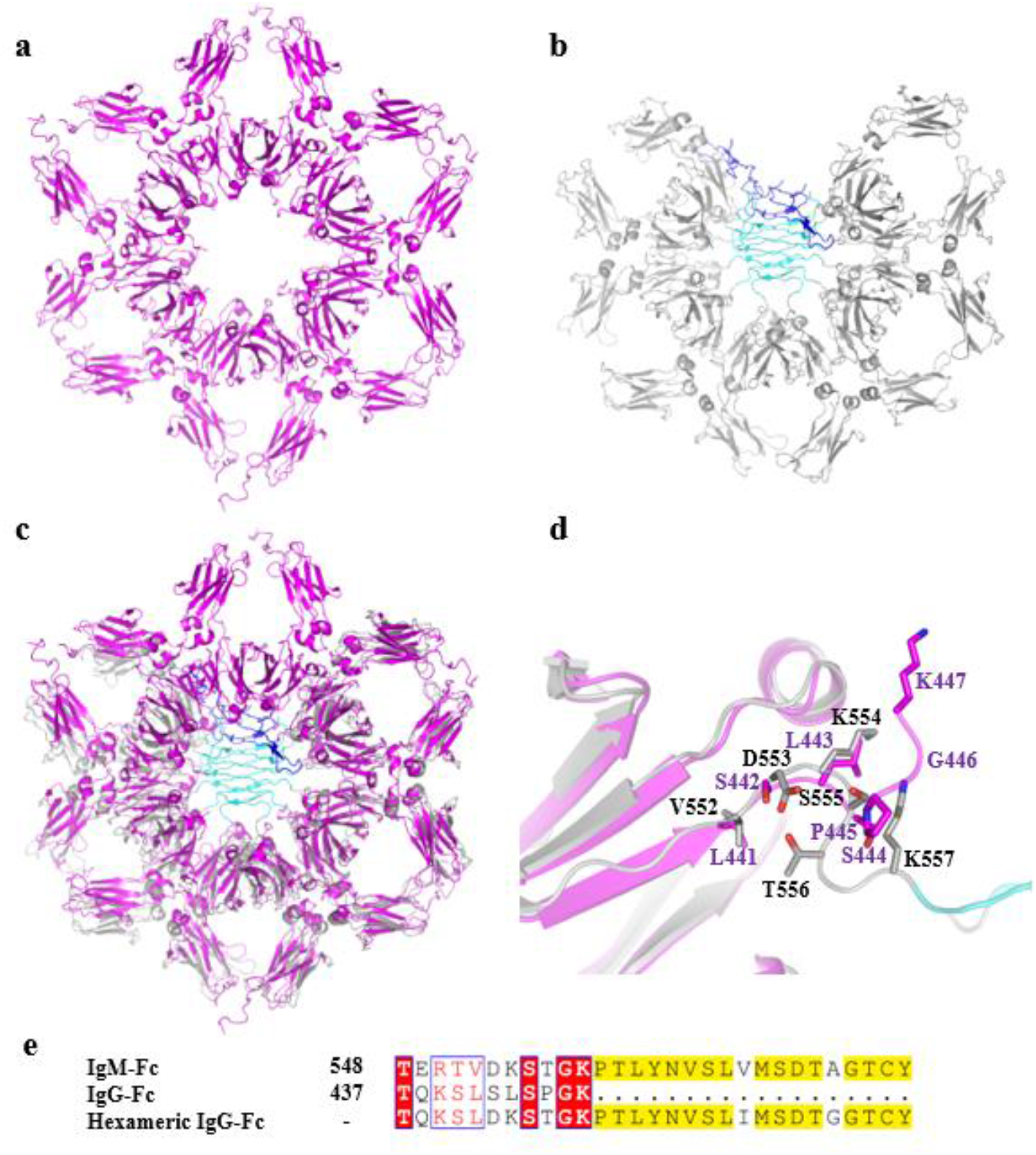
Engineering of hexameric IgG. **a,** IgG crystal packing formed IgG-Fc hexamer (purple, PDB code 1HZH). **b,** Cryo-EM structure of IgM core (PDB code 6KXS). IgM-Fcμ3-4, tailpiece and J-chain are colored in gray, cyan and blue respectively. **c,** Overlap of IgG-Fc hexamer and IgM core. **d,** Comparison of IgG-Fc and IgM-Fc C-terminal residues to show different orientations due to the difference of P445 in IgG-Fc and T556 in IgM-Fc. **e,** sequence alignment of IgG-Fc, IgM-Fc and engineered hexameric IgG-Fc C-terminal residues.

### Cryo-EM structure of Hexameric IgG-Fc

The flexible fab arms of IgG were removed during construct generation in order to obtain structure of the rigid IgG-Fc, which is the central region mediating polymerization of hexameric IgG^2^. This polymeric IgG-Fc was expressed in mammalian cells and purified for cryo-EM study as described in the Methods. Three-dimensional (3D) classification with Relion^11^ led to cryo-EM reconstruction of the hexameric IgG-Fc at nominal resolution of 3.7 Å (Fig. 2a and supplementary Fig. 2). The cryo-EM map allows model building of most regions. There is well-defined density for the interface between Fc4B and Fc5A (Supplementary Fig. 3), allowing for accurate modeling. Initial model was built by using the crystal packing formed polymeric IgG-Fc (PDB code 1HZH) and IgM tailpiece (PDB code 6KXS), and subsequent refinement was done in PHENIX^12^.

**Figure 2.**
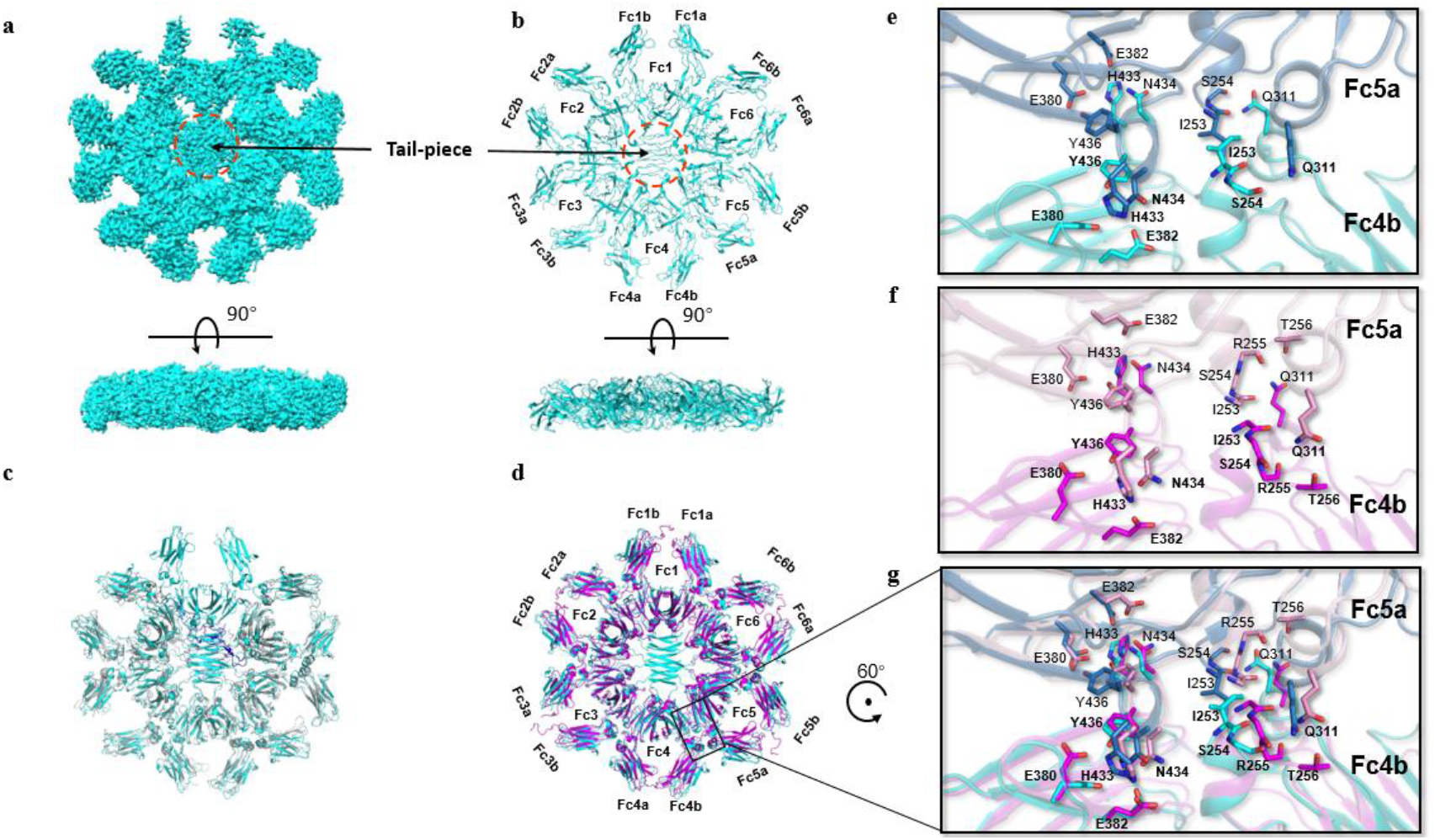
Cryo-EM structure of hexameric IgG. **a, b,** The cryo-EM density map and structure of IgG-Fc hexamer (cyan). **c, d,** Comparison of cryo-EM structure of IgG-Fc hexamer (cyan) with IgM or IgG crystal packing formed IgG-Fc hexamer (purple). IgM-Fcμ3-4, tailpiece and J-chain are colored in gray, light blue and blue respectively. **e-g,** Comparison of Fc-Fc interaction residues in cryo-EM structure of IgG-Fc hexamer (cyan and light blue) with IgG crystal packing formed IgG-Fc hexamer (purple and pink).

The IgG-Fc hexamer forms perfect hexagon symmetry with the six Fcs lie in a plane (Fig. 2b). Unlike pentameric IgM structure (Fig. 2c), this IgG-Fc hexamer is in agreement with crystal packing formed IgG-Fc hexamer (Fig. 2d). L309C of adjacent Fcs are close to each other, but do not form inter-Fc disulfide bonds to stabilize the hexamer as described previously^8^.

### Fc-Fc interaction in a mutual lock-and-key mode

The cryo-EM map includes well-defined features for Fc-Fc interactions composed of only 8 residues I253, S254, Q311, E380, E382, H433, N434, Y436 from each Fc (Fig. 2e). Q311, H433-N434, Y436 from Fc4b contact with I253-S254, E380 and E382, Y436 in Fc5a, respectively. These residues contribute a few of ionic interactions, hydrogen bonds and lack of hydrophobic interactions. This is highly consistent with crystal packing formed Fc-Fc interface, which has only two extra residues R255-T256 binding with Q311 (Fig. 2f–g).

Previous studies with Fc mutants at I253, H433 or N434 revealed a decreased binding with C1q and complement activation^7^. Of particular interest, side chain of residues I253, Q311, H433-N434 in one Fc protrude to three different cavities of neighboring Fc (Fig. 3a–b), and the reverse is also true. Therefore, residues forming the Fc-Fc interface contact in a mutual lock-and-key mode (Fig. 3c).

**Fig. 3.**
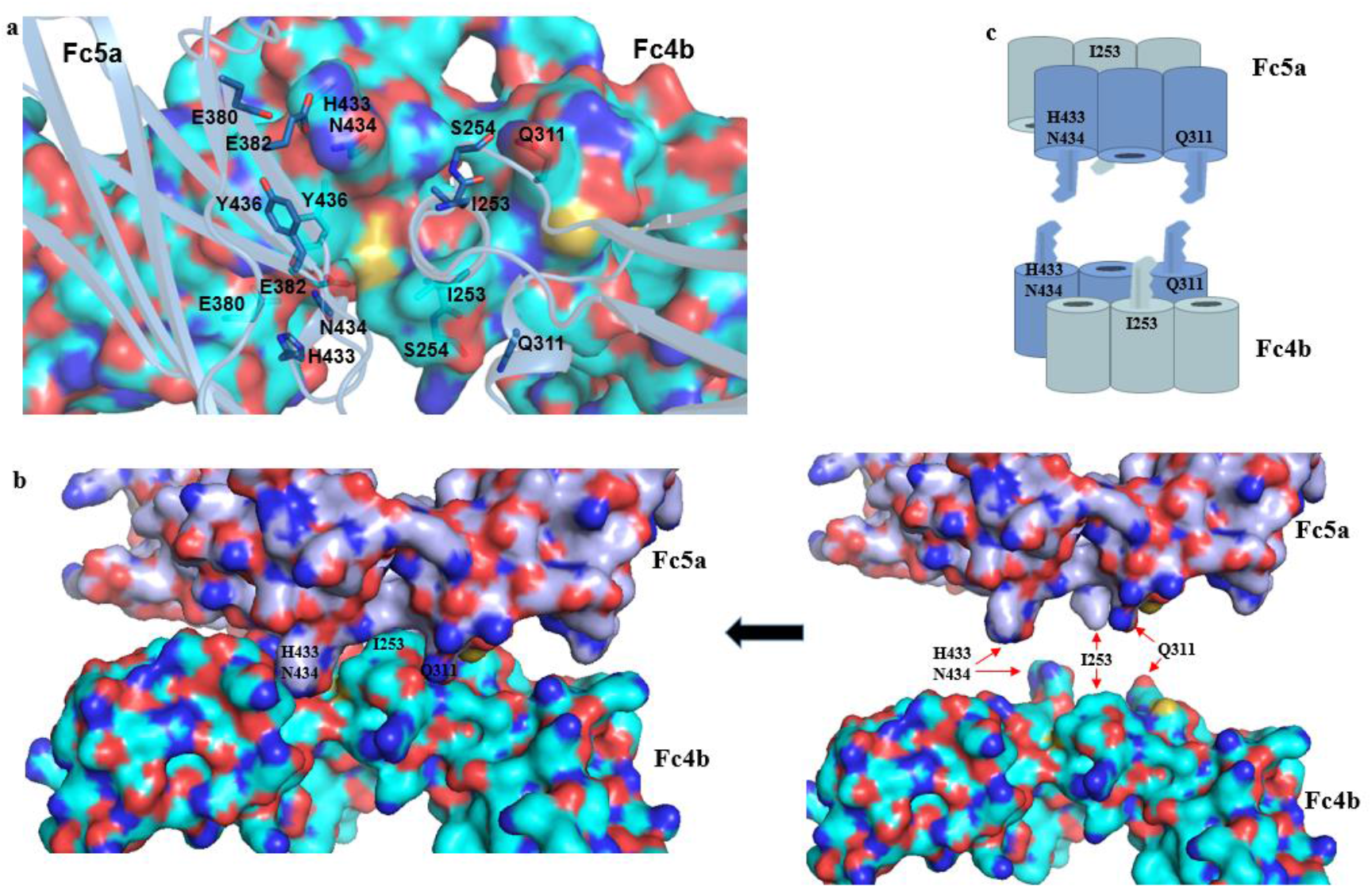
Fc-Fc interaction in a mutual lock-and-key mode. **a, b,** side chain of residues I253, Q311, H433-N434 in one Fc protrude to three different cavities of neighboring Fc. **c,** cartoon representation of the Fc-Fc interaction in a mutual lock-and-key mode.

## Discussion

For a long period, the Fc-Fc interaction of IgG was not clear. This engineered hexameric IgG-Fc driven by IgM tailpiece fusion (supplementary Fig. 4a) provides the first atomic structure of IgG-Fc hexamer and reveals a mutual lock-and-key mode of Fc-Fc interaction. IgG exists as monomer in solution due to the weak Fc-Fc interaction. However, crosslinking by Fcγ receptors on Fc N-terminus or antigens by Fab binding might also promote the formation of hexamer (supplementary Fig. 4a). Therefore, this weak interaction but rigid assembled IgG-Fc hexamer may play important roles in IgG function.

Previous low resolution map of IgG-C1 complex by cryo-ET shows evidence for IgG hexamer formation after antigen binding to activate complement system. In addition, cross-linking of antibody and antigens forms lattice-like network of immune complex and can be cleared via complement system^13^. Taken together, the IgG might exist as hexamer in immune complex formation and clearance (supplementary Fig. 4b).

Hexameric IgG via IgM tailpiece fusion or Fc-Fc interface mutations have been shown to increase CDC function, crosslinking ability of cell surface receptors or SARS-CoV-2 virus neutralization ability^14,15^. Our hexameric IgG-Fc structure provides detail information of the Fc-Fc interaction and guidance for further engineering for therapeutic application of IgG hexamer.

## Acknowledgements

This work was supported by the Innovent Biologics (Suzhou) Co., Ltd. We thank Shuimu BioSciences for providing the cryo-EM data collection support.

## Author Contributions

D.Y. designed the Hexameric IgG, conceived the experiments and wrote the manuscript with input from all authors. S.Z. led protein expression and purification. W.Y., X.F., H.N., Y.G., D.B., Z.S., Z.W. J.Z., L.L., J.S., N.Z., performed experiments. M.Y., Y.L., X.X., B.C., C.G. provided funding and critical support for the work.

