## supplementary materials3 for "Structure of IgG-Fc hexamer reveals a mutual lock-and-key mode of Fc-Fc interaction"

**This PDF file includes:**

Materials and Methods
References
Figs. S1 to S4
Tables S1

### Materials and Methods

#### Construction, protein expression and purification of hexameric IgG-Fc

The genes encoding signal peptide and IgG-Fc residues D221-L441 fused with residues D553-Y576 of IgM, including 3 point mutations, L309C, V567I and A572G (Supplementary Fig. 1), were synthesized and cloned into a modified pcDNA3.1 vector. This construct was used to express recombinant hexameric IgG-Fc in HEK293 cells (Innovent BiologicsCo., Ltd., Suzhou, China) by transient transfection. Then, we obtained the antibodies through standard methods using protein A chromatography. After purification, size-exclusion chromatography (SEC) and SDS-PAGE analyses demonstrated that recombinant antibodies exhibited high purity and existed in a homogeneous state.

#### Cryo-EM sample preparation and data acquisition

Samples of recombinant hexameric IgG-Fc at a concentration 1.93 mg/ml were applied for cryo-EM grids preparation using a Thermo Fisher Vitrobot Mark IV plunger. The recombinant hexameric IgG-Fc of volume 3.5 μL was placed on a glow discharged holey carbon grid (LeHua, Ni-Ti R1.2/1.3). Then excess solution from the grid was blotted for 2.5–3.0 s in 100% humidity at 8 °C before the grid was flash frozen in liquid ethane cooled at liquid-nitrogen temperature.

Cryo-EM data were collected on a Thermo Fisher TitanKrios G3i electron microscope equipped with a Gatan K3 direct electron counting camera. The microscope was operated at 300 kV, and images of the specimen were recorded with a defocus range from –1.1 to –1.9 μm at a calibrated magnification of 105,000, yielding to a pixel size of 0.85 Å. The movie stacks, each containing 50 frames, were recorded with the semi-automated low-dose acquisition program EPU, with a total dose of approximately 55 electrons/Å2.

**Image processing**

For data processing, all movie frames were motion-corrected using Relion3.1[1] and the contrast-transfer function (CTF) parameters were estimated by GCTF. A total of 1,052,204 particles were automatically picked from 2880 micrographs. After a series of operations containing particles selection, 2D classification, 3D classification performed using Relion3.1, a data of 300,561 particles were subjected to 3D auto-refinement without symmetry, reporting the global resolution at 3.7 Å. Reconstruction resolution were determined based on the gold-standard Fourier shell correlation (FSC =0.143). Half-reconstruction was used to determine the local resolution of each map.

**Model building and refinement**

The initial model of hexameric IgG-Fc was built by using the structure of IgG1 (Protein Data Bank accession (PDB):1HZH) and IgM (PDB: 6KXS). The models were docked into the cryo-EM density map, followed by iterative manual adjustment and real-space refinement using COOT[2] and PHENIX. Model overfitting was evaluated against one cryo-EM half map. The final refinement statistics of our model are shown in Supplementary information, Table S1.

1 Scheres, S. H. RELION: implementation of a Bayesian approach to cryo-EM structure determination. *J Struct Biol* **180**, 519-530, doi:10.1016/j.jsb.2012.09.006 (2012).

2 Emsley, P. & Cowtan, K. Coot: model-building tools for molecular graphics. *Acta Crystallogr D Biol Crystallogr* **60**, 2126-2132, doi:10.1107/S0907444904019158 (2004).

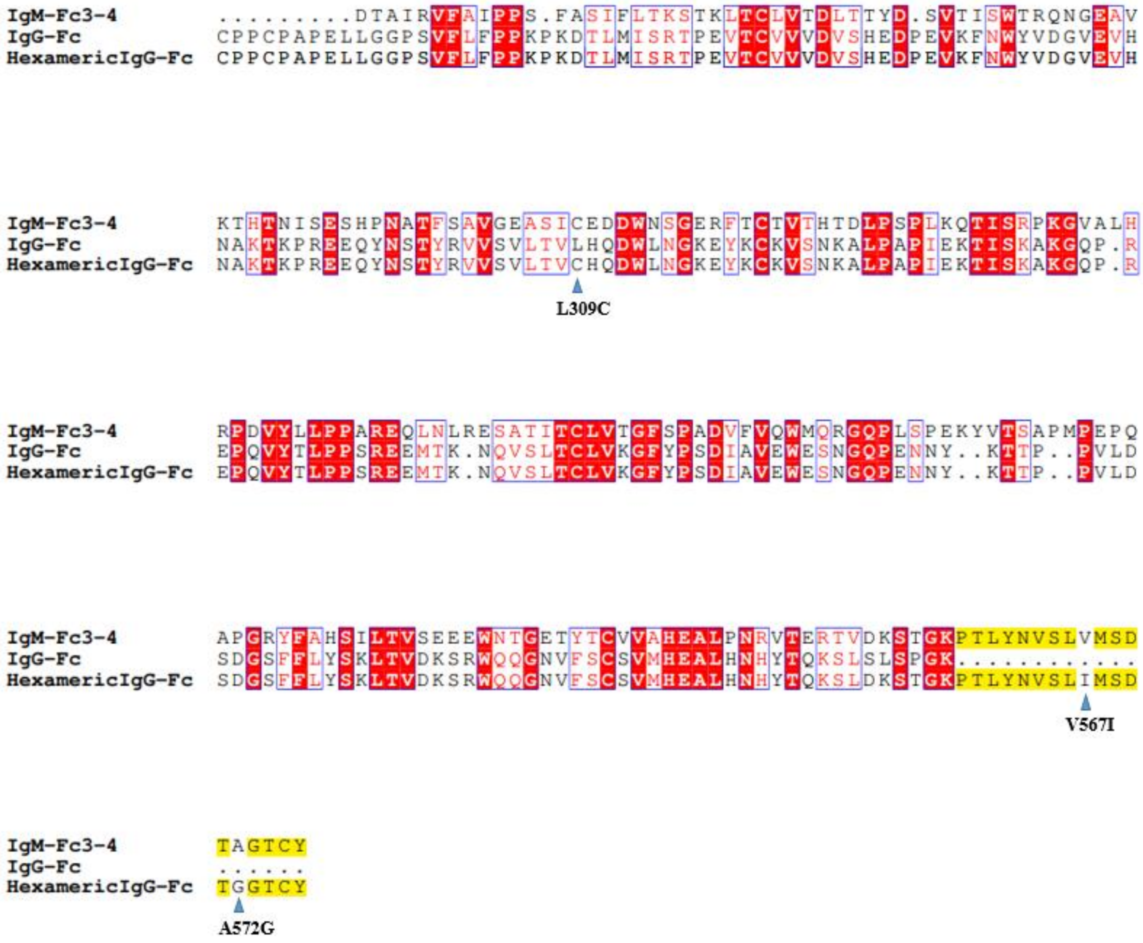

**Supplementary Fig.S1 | Sequence alignment of IgM-Fcμ3-4, IgG-Fcγ2-3 and hexameric IgG-Fc.** Hexameric IgG-Fc has residues D221-L441 of IgG fused with residues D553-Y576 of IgM, including 3 point mutations, L309C, V567I and A572G.

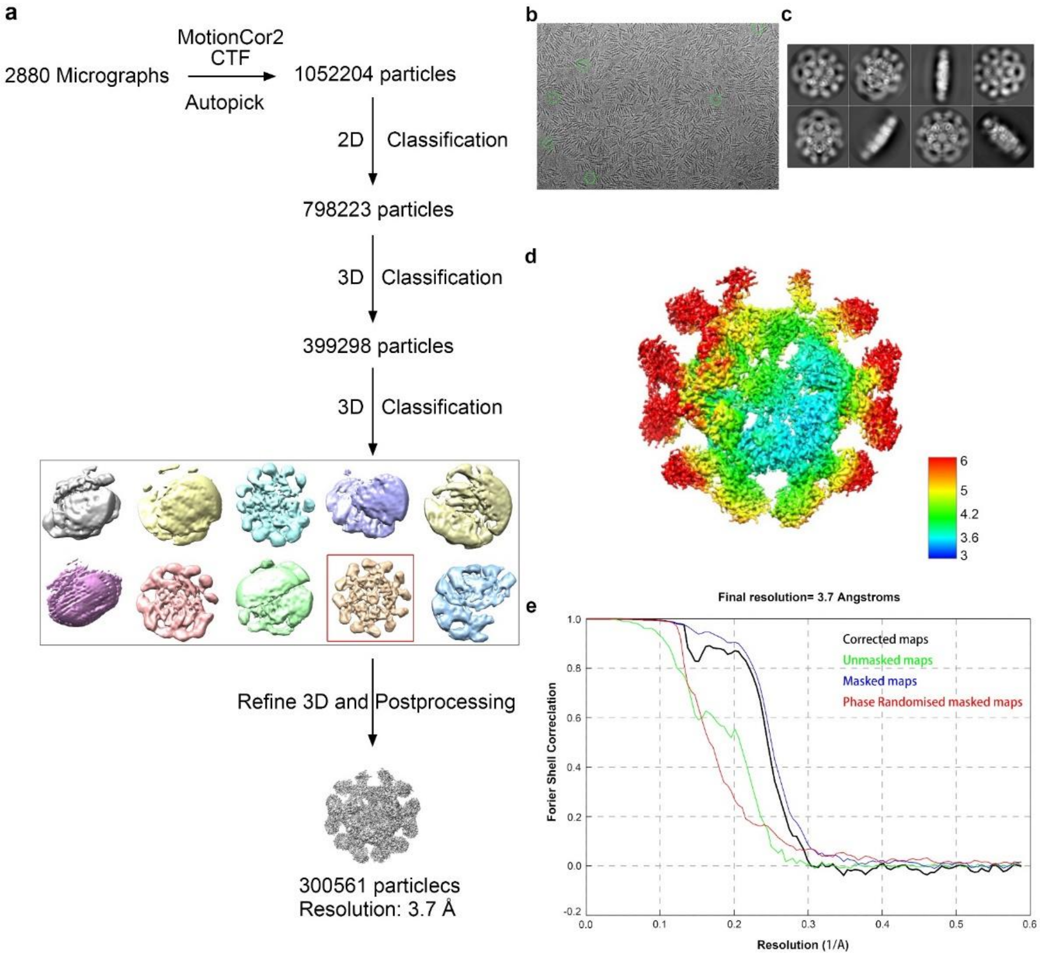

**Supplementary Fig.S2 | Cryo-EM data processing of the hexameric IgG-Fc. a,** Workflow for image processing. **b,c,** Representative image and 2D class averages of the protein showing different views. **d,** Local resolution of the hexameric IgG-Fc was shown in color-coded map. **e,** FSC curves of the final reconstruction showing the overall nominal resolution at 3.7 Å.

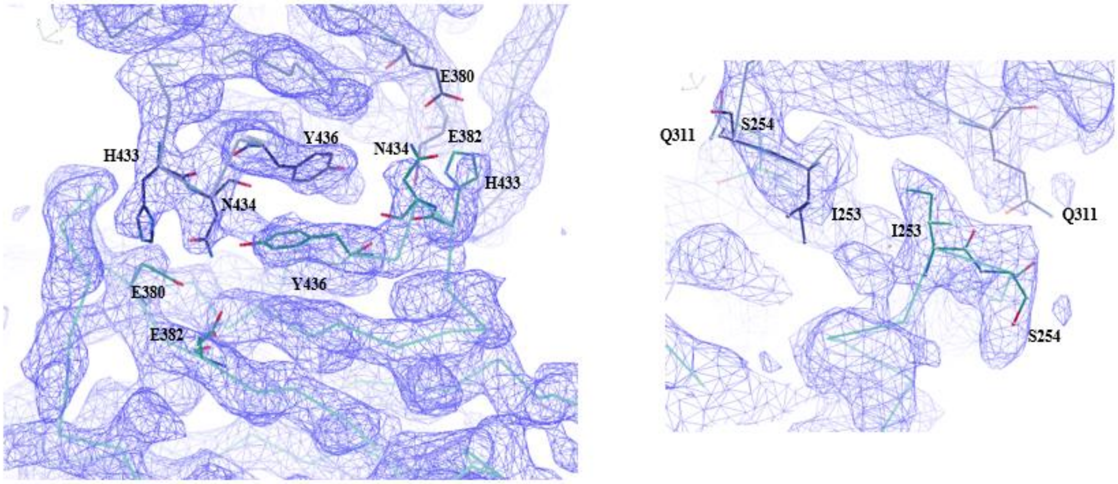

**Supplementary Fig.S3 | Cryo-EM map of key Fc-Fc contact residues in the hexameric IgG-Fc.**

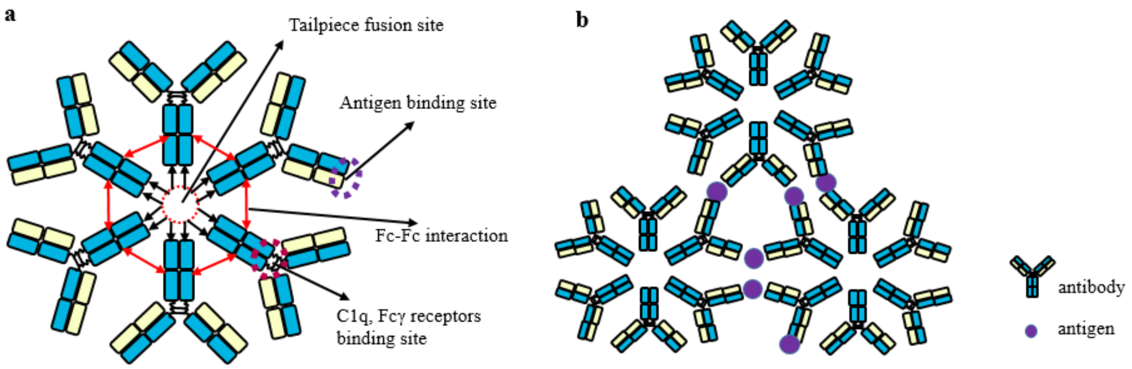

**Supplementary Fig.S4 | a,** cartoon representation of IgG hexamer showing tailpiece fusion site, antigen, C1q, Fc γ receptors binding sites and Fc-Fc interaction. **b,** cartoon representation of a model of immune complex.

| | IgG-Fc hexamer (EMDB-EMD-32933) (PDB-7X13) |
|---|---|
| **Data collection and processing** | |
| Magnification | 105,000 |
| Voltage (kV) | 300 |
| Electron exposure (e-Å$^2$) | 55 |
| Defocus range (μm) | -1.1 to -1.9 |
| Pixel size (Å) | 0.85 |
| Symmetry imposed | C1 |
| Initial particle images (no.) | 1052141 |
| Final particle images (no.) | 300561 |
| Map resolution (Å) | 3.7 |
| FSC threshold | 0.143 |
| Map resolution range (Å) | 3.0-6.0 |
| **Refinement** | |
| Initial model used (PDB code) | 6KXS and 1HZH |
| Model resolution (Å) | 3.7 |
| FSC threshold | 0.143 |
| Model resolution range (Å) | 3.0-6.0 |
| Map sharpening *B* factor (Å$^2$) | -181 |
| Model composition | |
| Non-hydrogen atoms | 21248 |
| Protein residues | 2668 |
| *B* factors (Å$^2$) | |
| Protein | 36.10 |
| R.m.s deviations | |
| Bond lengths (Å) | 0.002 |
| Bond angles (°) | 0.500 |
| Validation | |
| Molprobity score | 1.44 |
| Clashscore | 8.08 |
| Poor rotamers (%) | 0 |
| Ramachandran plot | |
| Favored (%) | 98.26 |
| Allowed (%) | 1.74 |
| Disallowed (%) | 0 |

**Supplementary Table 1 | Cryo-EM data collection, refinement and validation statistics.**
